# Pan-cancer imaging of TERT expression using deuterium magnetic resonance spectroscopy-based assessment of pyruvate metabolism

**DOI:** 10.1101/2021.10.05.462047

**Authors:** Georgios Batsios, Céline Taglang, Meryssa Tran, Nicholas Stevers, Carter Barger, Anne Marie Gillespie, Sabrina M Ronen, Joseph F Costello, Pavithra Viswanath

## Abstract

Telomerase reverse transcriptase (TERT) expression is indispensable for tumor immortality. Non-invasive methods of imaging TERT can, therefore, report on tumor proliferation and response to therapy. Here, we show that TERT expression is associated with elevated levels of the redox metabolite NADH in multiple cancers, including glioblastoma, oligodendroglioma, melanoma, neuroblastoma, and hepatocellular carcinoma. Mechanistically, TERT acts via the metabolic regulator FOXO1 to upregulate nicotinamide phosphoribosyl transferase, which is the key enzyme for NADH biosynthesis. Importantly, deuterium magnetic resonance spectroscopy (^2^H-MRS), which is a novel, clinically translatable metabolic imaging modality, can be leveraged for imaging TERT-linked NADH in preclinical tumor models *in vivo*. Since NADH is essential for pyruvate flux to lactate, ^2^H-MRS following administration of ^2^H-labeled pyruvate non-invasively visualizes TERT expression and reports on early response to therapy. Collectively, our study provides insights into the mechanisms of TERT-linked metabolic reprogramming and, importantly, establishes ^2^H-MRS as a pan-cancer strategy for imaging tumor immortality.

## INTRODUCTION

Progressive telomere shortening constitutes a natural barrier to cell proliferation^1^. Telomeres are cap-like structures composed of telomeric DNA and specialized proteins that protect the ends of linear chromosomes from damage during replication^1^. Telomerase reverse transcriptase (TERT) is the catalytic component of the enzyme telomerase that synthesizes telomeric DNA, and its expression is silenced in normal somatic cells, except for stem cells^1,2^. TERT expression is reactivated via mutations in the TERT promoter in ∼80% of human tumors, including glioblastomas, oligodendrogliomas, melanomas, hepatocellular carcinomas, and neuroblastomas^2,3^. TERT promoter mutations create a binding site for the transcription factor GABP, which then induces TERT transcription from the mutant TERT promoter^3,4^. Reactivation of TERT expression enables tumor cells to achieve immortality by overcoming telomere shortening and is an early and fundamental event in tumorigenesis^5-7^. Due to its essential role in sustaining proliferation, TERT is also a therapeutic target^1,2,8^. Inhibition of TERT function using small molecules such as 6-thio-2′-deoxyguanosine (6-thio-dG)^9,10^ or via antisense oligonucleotide-based inhibition of TERT expression^8^ have emerged as promising anti-cancer therapeutic strategies.

Magnetic resonance imaging (MRI) is a key tool for non-invasive diagnosis, staging and treatment response assessment in cancer^11^. However, anatomical imaging methods do not report on biological events that sustain tumor proliferation and can fail to distinguish tumor from surrounding normal tissue^12^. Importantly, changes in tumor size can be slow to occur following treatment, leading to incorrect assessments of disease progression^13-15^. Since TERT expression is specifically reactivated in cancer cells and plays a key role in sustained tumor proliferation^2^, TERT is a molecular biomarker of tumor immortality. Identifying MR-based biomarkers of TERT expression can, therefore, enable the non-invasive assessment of tumor proliferation and response to therapy.

Previous studies have linked TERT to redox homeostasis in cancer. TERT expression has been shown to alleviate oxidative stress via upregulation of reduced glutathione (GSH) in HeLa cells^16^. We and others have also shown that TERT expression in gliomas is associated with elevated levels of GSH, NADPH and NADH, which play key roles in redox homeostasis^17-19^. While these studies point to an association between TERT and altered redox in cancer, the precise mechanisms and the universality of TERT-linked metabolic reprogramming across multiple cancer types remain elusive.

Magnetic resonance spectroscopy (MRS) is a non-invasive, non-radioactive, MR-based method of interrogating metabolic activity in live cells, animals, and patients^13,20^. ^1^H-MRS examines the nuclear magnetic resonance of protons in cellular metabolites and provides a readout of steady-state metabolite levels^20^. Steady-state metabolite levels, however, do not accurately reflect the activity of a metabolic pathway and may not be sensitive to fluctuations in concentrations of cofactors such as NADH^21^. Recently, ^2^H-MRS, in which the nuclear magnetic resonance of the ^2^H moiety is interrogated, emerged as a versatile, innovative method of examining metabolic pathway activity *in vivo*^22^. Following administration of ^2^H-labeled substrates such as glucose, acetate or fumarate, flux to metabolic products can be quantified in real-time in a non-invasive manner in preclinical tumor models *in vivo*^23-27^. Importantly, the feasibility of ^2^H-MRS has been established in glioblastoma patients^23^, underscoring the clinical translatability of this method.

The goal of this study was to delineate the mechanisms linking TERT to redox across multiple cancers that reactivate TERT expression and to leverage this information for non-invasive ^2^H-MRS-based metabolic imaging of tumor burden and treatment response. We show that TERT inhibits the transcription factor FOXO1 in patient-derived models of glioblastomas, oligodendrogliomas, melanomas, neuroblastomas, and hepatocellular carcinomas. FOXO1, in turn, inhibits expression of nicotinamide phosphoribosyl transferase, the rate-limiting enzyme for NADH biosynthesis^28^. As a result, TERT expression results in elevated NADH. Importantly, we demonstrate that TERT-linked NADH can be visualized *in vivo* using ^2^H-MRS following administration of [U-^2^H]-pyruvate. Furthermore, imaging TERT using [U-^2^H]-pyruvate reports on response to chemotherapy and targeted TERT inhibitors at early timepoints preceding MRI-detectable alterations in tumor volume. Collectively, our study provides novel mechanistic insights into TERT biology and identifies an innovative, pan-cancer method for imaging TERT that can be utilized for *in vivo* assessment of tumor burden and response to therapy.

## RESULTS

### TERT expression is associated with elevated steady-state levels of NADH in cancer

The redox environment of mammalian cells is regulated by levels of redox-reactive small molecules including GSH, oxidized glutathione (GSSG), NADPH, NADP+, NADH and NAD+^29^. To determine whether TERT expression is linked to redox across multiple cancer types, we examined the effect of silencing TERT expression by RNA interference in preclinical models of glioblastoma (GBM1, GBM6), oligodendrogliomas (SF10417, BT88), melanoma (A375), neuroblastoma (SK-N-SH) and hepatocellular carcinoma (HepG2). We confirmed that TERT mRNA and telomerase activity were significantly reduced in TERT-cells relative to TERT+ in all our models (Fig. S1A-S1B). TERT silencing resulted in a significant drop in levels of both NADH and NAD+ in our models, leading to a drop in the NADH/NAD+ ratio (Fig. 1A-1C). In contrast, silencing TERT caused a significant reduction in GSH and NADPH in glioblastoma, oligodendroglioma and melanoma, but not in neuroblastoma or hepatocellular carcinoma (Fig. 1D-1E). There was no change in GSSG or NADP+ following TERT silencing in any of our models (Fig. S1C-S1D). These results identify elevated NADH and NAD+ as common metabolic alterations linked to TERT expression across multiple cancers^30^.

**Figure 1.**
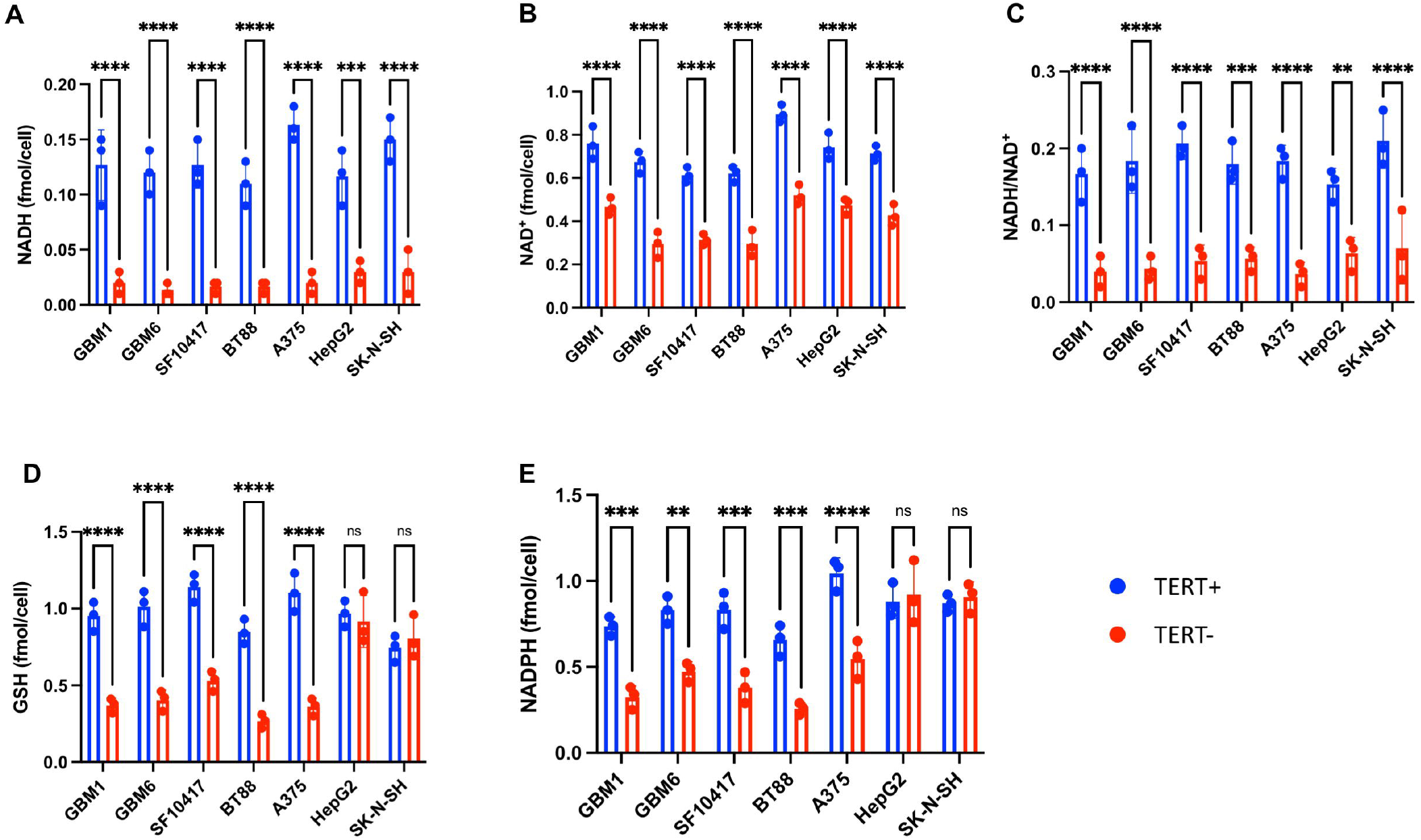
TERT expression is linked to levels of NADH and NAD^+^ in cancer cells. NADH **(A)**, NAD^+^ **(B)**, NADH/NAD^+^ ratio **(C)**, GSH **(D)** and NADPH **(E)** in TERT+ and TERT-glioblastoma (GBM1, GBM6), oligodendroglioma (SF10417, BT88), melanoma (A375), hepatocellular carcinoma (HepG2) and neuroblastoma (SK-N-SH) cells. Bars depict mean values and error bars represent standard deviation. * represents p<0.05, ** represents p<0.01, *** represents p<0.001 and **** represents p<0.0001.

### TERT elevates levels of NADH via upregulation of nicotinamide phosphoribosyl transferase (NAMPT)

Although NAD+ can be synthesized *de novo* from dietary tryptophan or nicotinic acid, most NAD+ in mammalian cells is recycled via the salvage pathway from nicotinamide (NAM), which is subsequently converted to nicotinamide mononucleotide (NMN) in a rate-limiting reaction catalyzed by NAMPT (see schematic in Fig. 2A)^28,30^. NMN is subsequently adenylated to NAD+. Silencing TERT significantly reduced NAMPT expression and activity in all our models (Fig. 2B-2C). Exogenous supplementation with NMN, which is the product of NAMPT, restored levels of NAD+ and NADH in TERT-cells to levels observed in TERT+ cells (Fig. 2D-2E), further confirming a role for NAMPT in TERT-mediated upregulation of NAD+ and NADH. In contrast, supplementation with tryptophan or nicotinic acid, which are precursors for *de novo* NAD+ biosynthesis^28,30^, did not rescue the reduction in NAD+ and NADH levels in TERT-cells (Fig. S2A-S2D). These results identify NAMPT as a molecular determinant of elevated NAD+ and NADH in TERT+ cells.

**Figure 2.**
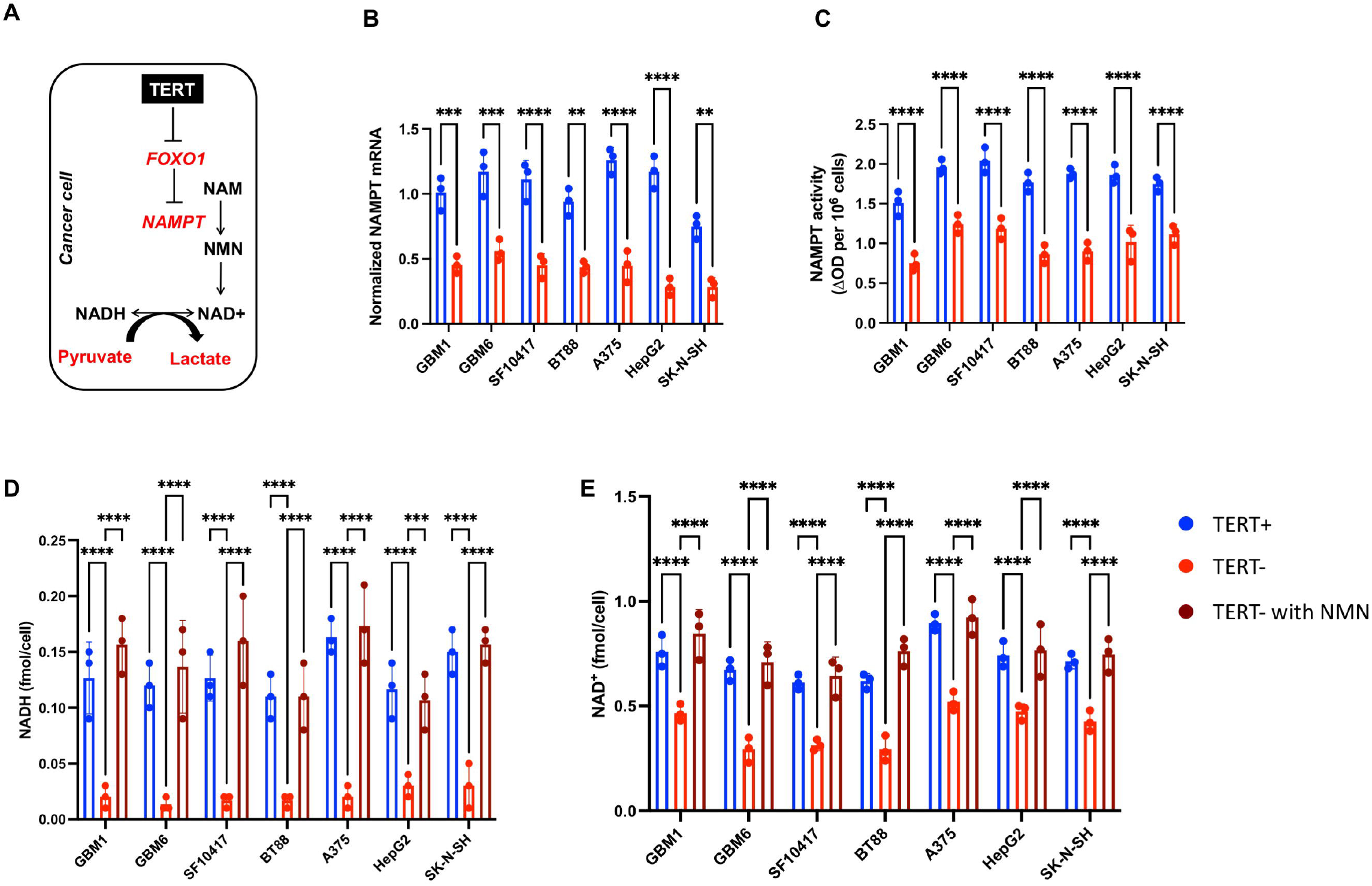
TERT elevates NADH and NAD^+^ via upregulation of NAMPT. **(A)** Schematic illustration of the link between TERT, FOXO1, NAMPT, NADH and pyruvate flux to lactate, which is dependent on concomitant conversion of NADH to NAD^+^. NAMPT mRNA **(B)** and activity **(C)** in TERT+ and TERT-glioblastoma (GBM1, GBM6), oligodendroglioma (SF10417, BT88), melanoma (A375), hepatocellular carcinoma (HepG2) and neuroblastoma (SK-N-SH) cells. Levels of NADH **(D)** and NAD^+^ **(E)** in TERT+ cells, TERT-cells and TERT-cells supplemented with NMN in glioblastoma (GBM1, GBM6), oligodendroglioma (SF10417, BT88), melanoma (A375), hepatocellular carcinoma (HepG2) and neuroblastoma (SK-N-SH) models. Bars depict mean values and error bars represent standard deviation. * represents p<0.05, ** represents p<0.01, *** represents p<0.001 and **** represents p<0.0001.

### TERT acts via FOXO1 to upregulate NAMPT expression and activity

The forkhead box O (FOXO) family of transcription factors, including FOXO1, play significant roles in regulating expression of genes involved in metabolism and redox homeostasis^31^. A major mechanism of regulation of FOXO1 activity in cancer cells involves inhibitory phosphorylation on 3 conserved serine/threonine residues (T24, S256, and S319)^31,32^. Phospho-FOXO1 is sequestered in the cytoplasm and excluded from the nucleus, which prevents transactivation of FOXO1 target genes in the nucleus^31,32^. To determine whether FOXO1 plays a role in TERT-induced upregulation of NAMPT, NAD+ and NADH, we first examined the effect of silencing TERT on FOXO1 phosphorylation and activity. As shown in Fig. 3A, levels of phospho-FOXO1 were significantly reduced in TERT-cells relative to TERT+ in our glioblastoma (GBM1), oligodendroglioma (BT88), melanoma (A375), neuroblastoma (SK-N-SH) and hepatocellular carcinoma (HepG2) models. There was no difference in total FOXO1 between TERT+ and TERT-cells (Fig. 3A). Concomitantly, FOXO1 transcription factor activity, as quantified by an ELISA assay that detects FOXO1 in its active DNA-bound form, was elevated in TERT-cells relative to TERT+ (Fig. 3B) in all our models. These results suggest that TERT expression is associated with inactivation of FOXO1 via inhibitory phosphorylation.

**Figure 3.**
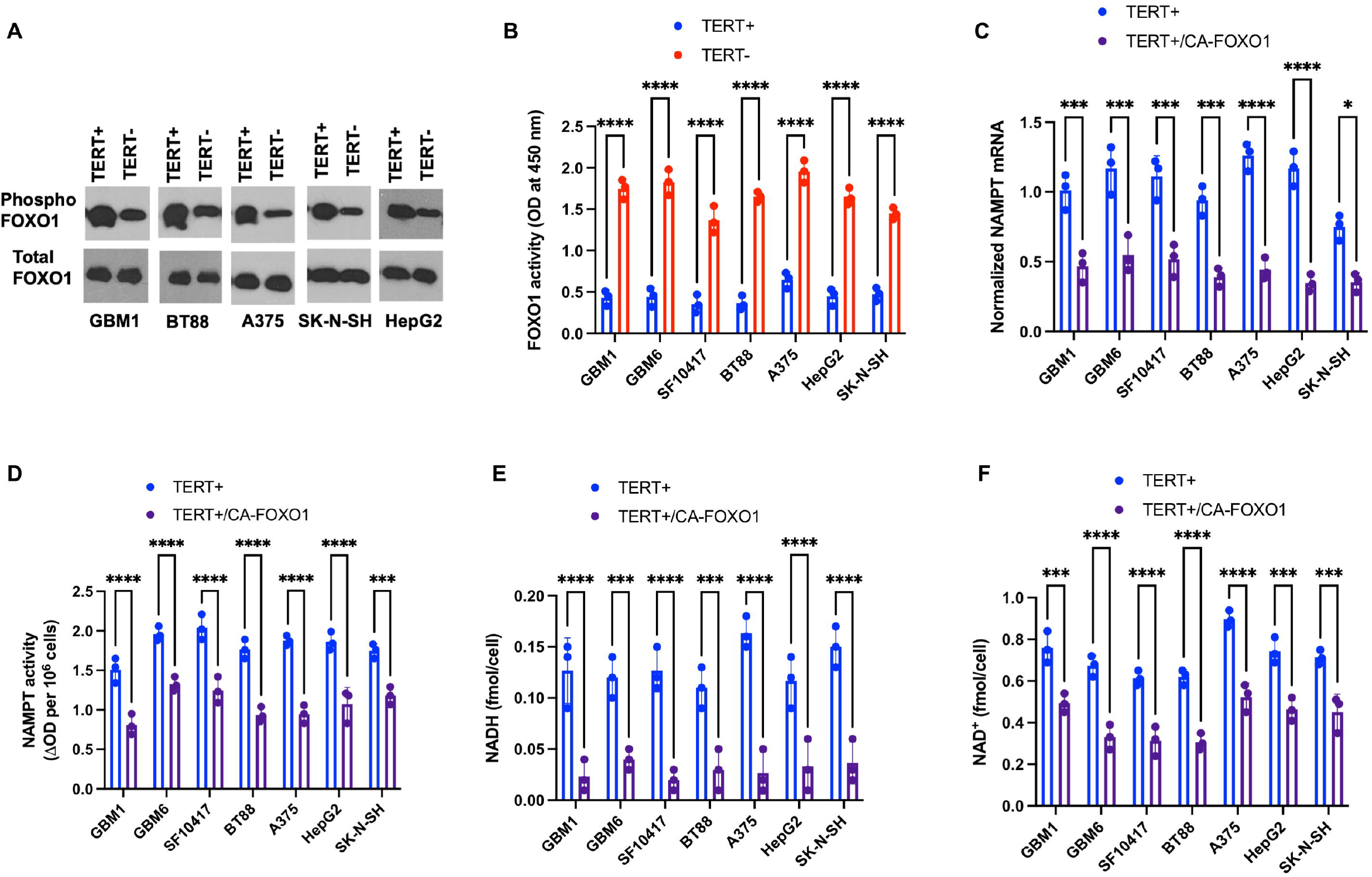
TERT acts via the FOXO1 transcription factor to upregulate NAMPT and NADH. **(A)** Western blots for phosphorylated and total FOXO1 protein in TERT+ and TERT-glioblastoma (GBM1), oligodendroglioma (BT88), melanoma (A375), neuroblastoma (SK-N-SH) and hepatocellular carcinoma (HepG2) cells. **(B)** FOXO1 transcription factor activity in TERT+ and TERT-glioblastoma (GBM1, GBM6), oligodendroglioma (SF10417, BT88), melanoma (A375), hepatocellular carcinoma (HepG2) and neuroblastoma (SK-N-SH) cells. Effect of expressing a constitutively active form of FOXO1 (CA-FOXO1) in TERT+ cells on NAMPT mRNA **(C)**, NAMPT activity **(D)**, NADH **(E)** and NAD^+^ **(F)** in glioblastoma (GBM1, GBM6), oligodendroglioma (SF10417, BT88), melanoma (A375), hepatocellular carcinoma (HepG2) and neuroblastoma (SK-N-SH) models. Bars depict mean values and error bars represent standard deviation. * represents p<0.05, ** represents p<0.01, *** represents p<0.001 and **** represents p<0.0001.

We then exogenously expressed a constitutively active form of FOXO1 (hereafter named CA-FOXO1) in TERT+ cells for all our cancer models^32^. CA-FOXO1 is a FLAG-tagged form of FOXO1 in which all three phosphorylation sites have been mutated to alanine (T24A, S256A, and S319A) and which is, therefore, constitutively active^32^. We confirmed CA-FOXO1 expression in transfected TERT+ cells via western blotting for the FLAG tag (Fig. S3A) and by confirming elevated FOXO1 transcription factor activity (Fig. S3B). As shown in Fig. 3C-3F, expression of CA-FOXO1 in TERT+ cells mimicked TERT silencing and caused a significant reduction in NAMPT expression and activity as well as levels of NAD+ and NADH. Collectively, the results presented in this section suggest that NAMPT expression and activity are negatively regulated by FOXO1 and that inhibitory phosphorylation of FOXO1 by TERT results in elevated NAMPT activity and consequently elevated NAD+ and NADH levels (refer to schematic summary in Fig. 2A).

### [U-^2^H]-pyruvate can non-invasively monitor TERT expression in preclinical tumor models ***in vivo***

NADH is an essential cofactor in the conversion of pyruvate to lactate and studies have demonstrated that lactate production from pyruvate reflects intracellular NADH^33^ and the NADH/NAD+ ratio^34^. Since our data points to elevated NADH and NADH/NAD+ ratios in TERT+ cells, we questioned whether lactate production from pyruvate is altered upon TERT silencing in our tumor models. To this end, we used ^2^H-MRS to monitor the conversion of [U-^2^H]-pyruvate to [U-^2^H]-lactate in live cells. Representative ^2^H-MR spectra acquired from neat cell culture medium (prior to incubation with cells) showed peaks for the natural abundance of semi-heavy water (HDO, 4.75 ppm) and [U-^2^H]-pyruvate (2.4 ppm; Fig. 4A). ^2^H-MR spectra acquired following incubation of live BT88 cells with media containing [U-^2^H]-pyruvate showed clear evidence of lactate (1.3 ppm) production with higher levels in BT88 TERT+ cells relative to TERT-cells (Fig. 4A). Importantly, quantification of the results across our TERT+ tumor models indicated that silencing TERT significantly reduced lactate production from [U-^2^H]-pyruvate, consistent with reduced levels of NADH upon TERT silencing (Fig. 4B).

**Figure 4.**
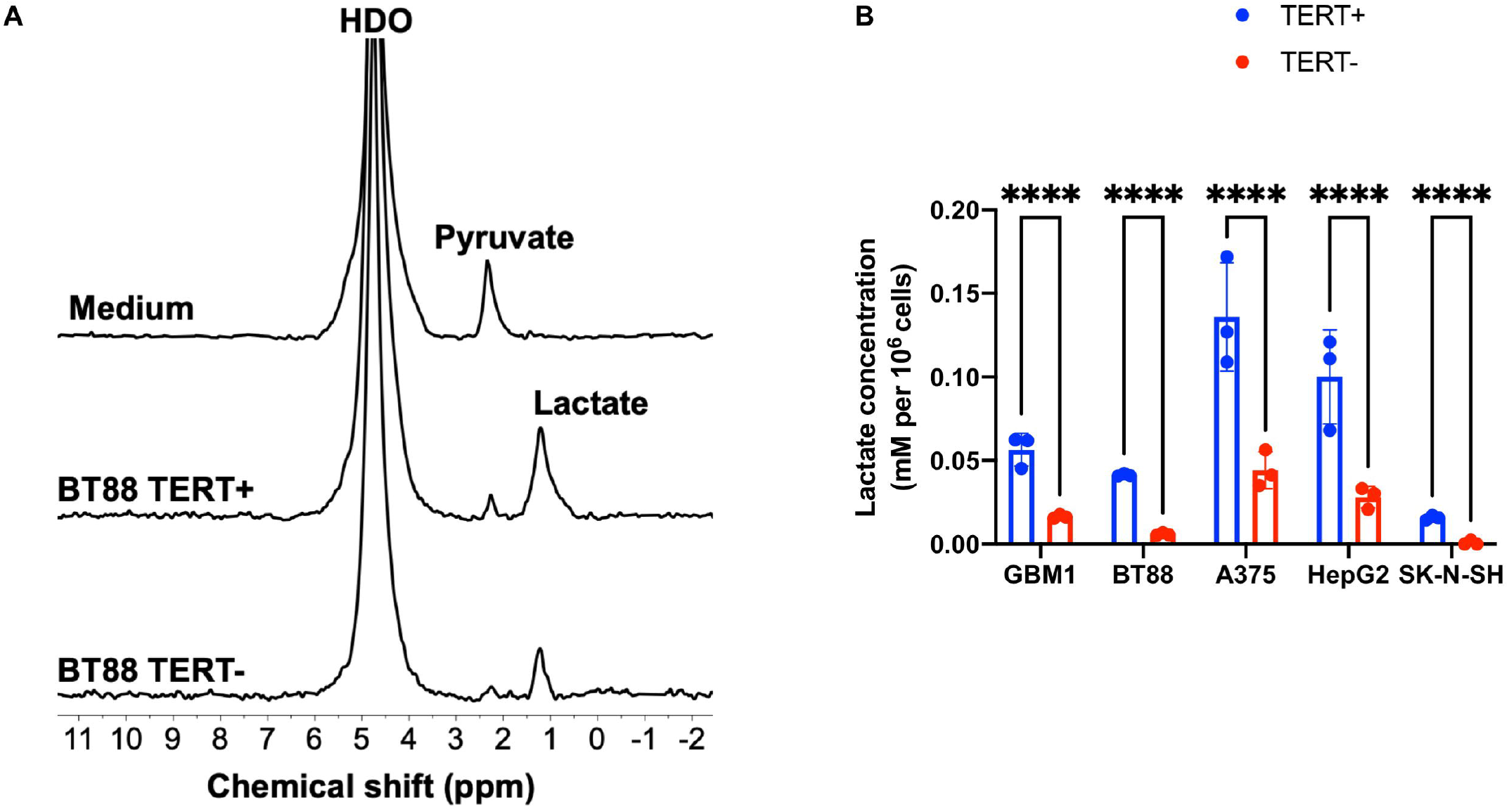
[U-^2^H]-pyruvate can non-invasively monitor TERT expression in live cells. **(A)** Representative ^2^H-MR spectra from TERT+ and TERT-BT88 cells cultured in medium containing 10 mM [U-^2^H]-pyruvate for 72 h. A representative spectrum acquired from neat culture medium containing 10 mM [U-^2^H]-pyruvate is also shown. Peaks for semi-heavy water (HDO; 4.75 ppm), pyruvate (2.4 ppm) and lactate (1.3 ppm) are labeled. **(B)** Effect of TERT silencing on ^2^H-lactate production from [U-^2^H]-pyruvate in glioblastoma (GBM1), oligodendroglioma (BT88), melanoma (A375), hepatocellular carcinoma (HepG2) and neuroblastoma (SK-N-SH) cells. Bars depict mean values and error bars represent standard deviation. * represents p<0.05, ** represents p<0.01, *** represents p<0.001 and **** represents p<0.0001.

We then examined the ability of [U-^2^H]-pyruvate to monitor TERT expression *in vivo*. To begin with, we examined [U-^2^H]-pyruvate metabolism in HepG2 tumors in which TERT expression has been silenced in a doxycycline-inducible manner. We first confirmed that TERT expression and telomerase activity were significantly reduced by doxycycline treatment in HepG2_dox-TERT_ cells (Fig. S4A-S4B). Doxycycline-induced TERT silencing resulted in a significant reduction in levels of NADH and NAD+ and a significant increase in FOXO1 transcription factor activity (Fig. 5A-5C). We then examined mice bearing subcutaneous HepG2_dox-TERT_ tumors that were treated with vehicle-control (saline) or doxycycline. TERT expression and telomerase activity were significantly reduced at day 8 relative to day 0 in doxycycline-treated, but not saline-treated tumor tissues (Fig. S4C-S4D). Following intravenous injection of a bolus of [U-^2^H]-pyruvate, metabolism to lactate was quantified prior to (day 0) and after (day 7) treatment with vehicle-control or doxycycline. A representative array of non-localized ^2^H-MR spectra acquired following intravenous injection of a bolus of [U-^2^H]-pyruvate in a HepG2_dox-TERT_ tumor-bearing mouse is shown in Fig. 5D. Prior to injection, only the HDO peak was observed. Following administration of [U-^2^H]-pyruvate, the spectral array showed dynamic production of lactate (Fig. 5D). We detected a small peak for pyruvate in the spectral array, which presumably reflects the limited endogenous pyruvate pool (∼0.2 mM)^35,36^. As expected, there was no significant difference in lactate concentration at day 0 between saline- and doxycycline-treated mice (Fig. 5E). Lactate production from [U-^2^H]-pyruvate was significantly reduced at day 7 relative to day 0 in doxycycline-treated mice, while, in contrast, lactate production was significantly higher at day 7 relative to day 0 in saline-treated animals (Fig. 5E). Importantly, assessment of tumor volume by T1-weighted MRI indicated that there was no significant difference in tumor volume between day 0 and day 7 in doxycycline-treated animals, consistent with prior studies indicating that there is a lag period before the onset of cell death following loss of TERT expression^37^ (5F-5G). These results indicate that metabolic imaging using [U-^2^H]-pyruvate reports on inhibition of TERT expression *in vivo* prior to MRI-detectable volumetric alterations.

**Figure 5.**
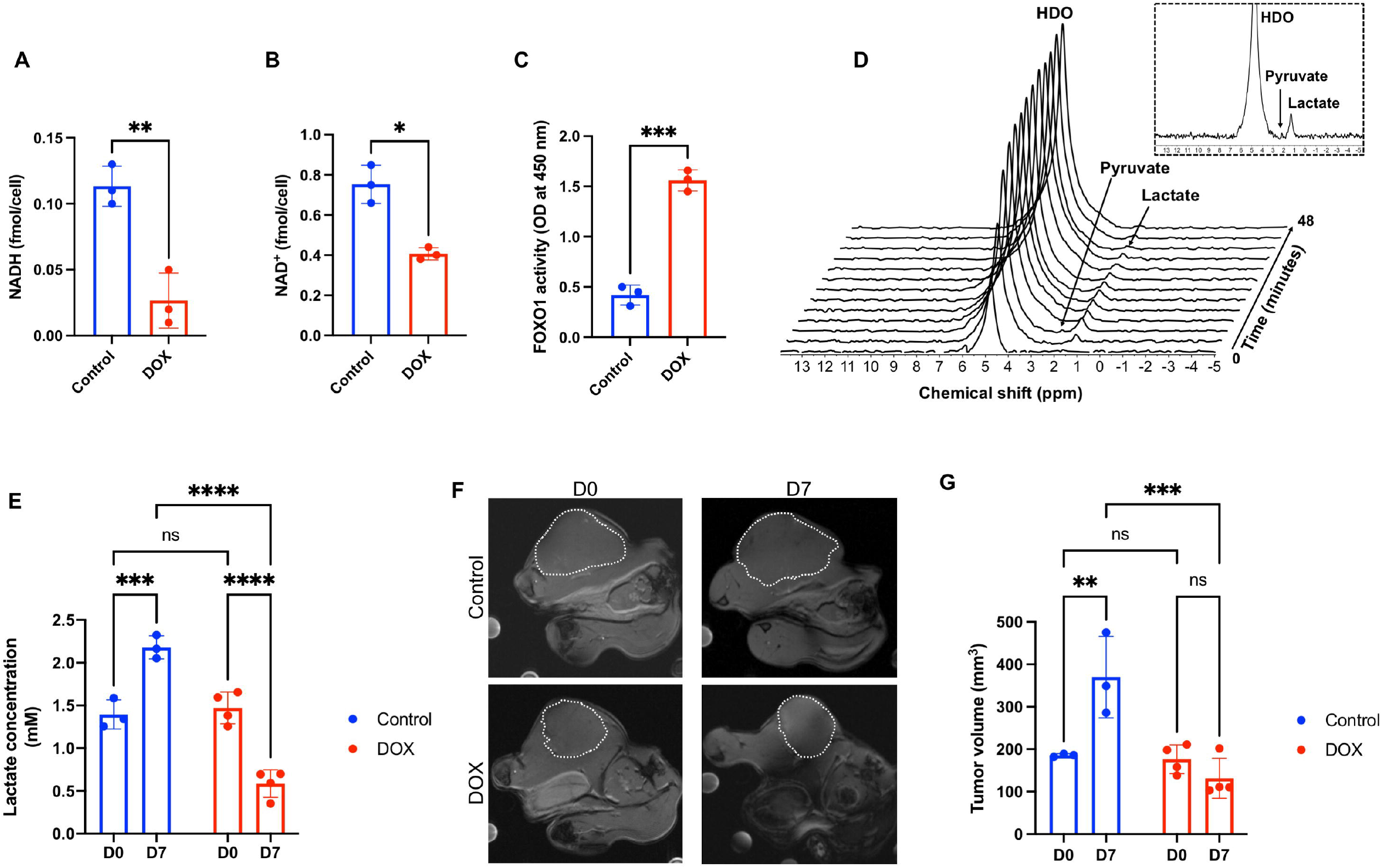
[U-^2^H]-pyruvate can non-invasively monitor TERT expression *in vivo*. Effect of silencing TERT via doxycycline treatment on NADH **(A)**, NAD^+^ **(B)** and FOXO1 transcription factor activity **(C)** in HepG2_dox-TERT_ cells. **(D)** Representative ^2^H-MR spectral array acquired from a saline-treated mouse bearing a subcutaneous HepG2_dox-TERT_ tumor following intravenous injection of [U-^2^H]-pyruvate. The first spectrum is acquired prior to [U-^2^H]-pyruvate injection. Inset shows an expansion of the spectrum with the highest lactate signal (2^nd^ spectrum following [U-^2^H]-pyruvate injection). **(E)** Concentration of ^2^H-lactate produced from [U-^2^H]-pyruvate on day 0 and day 7 in saline- and doxycycline-treated mice bearing subcutaneous HepG2_dox-TERT_ tumors. **(F)** Representative T1-weighted MRI at day 0 and day 7 of saline-treated (top row) and doxycycline-treated (bottom row) mice bearing subcutaneous HepG2_dox-TERT_ tumors. **(G)** Tumor volume at day 0 and day 7 in saline- and doxycycline-treated mice bearing subcutaneous HepG2_dox-TERT_ tumors. Bars depict mean values and error bars represent standard deviation. * represents p<0.05, ** represents p<0.01, *** represents p<0.001 and **** represents p<0.0001.

Since TERT expression is a hallmark of tumor immortality^1,2^, imaging TERT can enable assessment of tumor burden *in vivo*. We, therefore, examined the ability of [U-^2^H]-pyruvate to differentiate between TERT+ tumor and TERT-normal brain in mice bearing orthotopic glioblastoma (GBM1, GBM6) and oligodendroglioma (SF10417, BT88) tumor xenografts. Fig. 6A shows representative spectra from non-localized ^2^H-MRS studies in a mouse bearing an orthotopic SF10417 tumor vs. a tumor-free healthy control. Quantification of the data indicated that lactate production from [U-^2^H]-pyruvate was significantly higher in mice bearing orthotopic SF10417, BT88, GBM1 and GBM6 tumors relative to healthy controls (Fig. 6B). To further confirm these results and examine the spatial distribution of [U-^2^H]-pyruvate metabolism *in vivo*, we performed 2D chemical shift imaging (CSI) studies in mice bearing orthotopic GBM1 and BT88 tumor xenografts. Representative spectra from tumor and contralateral normal brain voxels in a mouse bearing an orthotopic GBM1 tumor xenograft are shown in Fig. 6C. Importantly, examination of the spatial distribution of the lactate signal overlaid over the corresponding anatomical MRI indicated that lactate production from [U-^2^H]-pyruvate was able to delineate tumor from surrounding contralateral normal brain in both the BT88 oligodendroglioma (Fig. 6D) and GBM1 glioblastoma (Fig. 6E) models. Quantification of the data confirmed significantly higher levels of lactate in tumor vs. contralateral heathy brain in both BT88 and GBM1 models (Fig. 6F). Collectively, the results presented in this section indicate that lactate production from [U-^2^H]-pyruvate is a non-invasive biomarker of TERT expression *in vivo* and point to its utility for *in vivo* imaging of tumor burden in preclinical cancer models.

**Figure 6.**
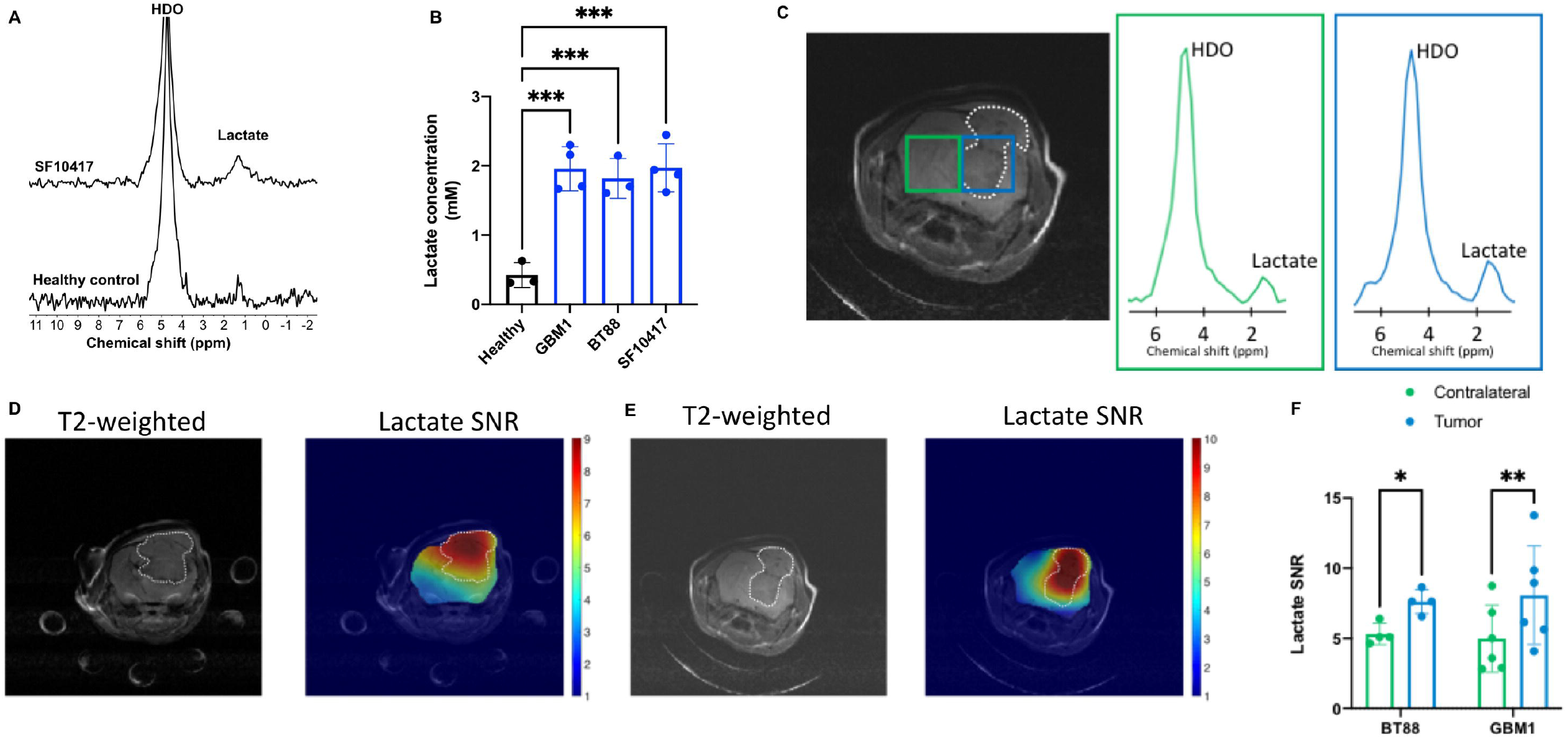
[U-^2^H]-pyruvate metabolism to lactate differentiates tumor from normal brain in patient-derived glioma models *in vivo*. **(A)** Representative ^2^H-MR spectra acquired after intravenous injection of [U-^2^H]-pyruvate into a mouse bearing an orthotopic SF10417 oligodendroglioma tumor (top row) or a tumor-free healthy control (bottom row). In each case, the spectra with the maximum lactate signal (2^nd^ spectrum following injection of [U-^2^H]-pyruvate] are shown. **(B)** Quantification of ^2^H-lactate production from [U-^2^H]-pyruvate in mice carrying orthotopic GBM1 glioblastoma tumors, orthotopic BT88 oligodendroglioma tumors or tumor-free healthy mice. **(C)** Representative 2D CSI data showing the T2-weighted MRI and corresponding ^2^H-MR spectra from voxels placed over the tumor and contralateral normal brain in a mouse bearing an orthotopic GBM1 tumor xenograft. Tumor voxel and spectra are shown in blue and contralateral normal brain voxel and spectra in green. The tumor region is contoured by white dotted lines. Representative heatmaps of ^2^H-lactate SNR from 2D CSI studies in mice bearing orthotopic BT88 **(D)** or GBM1 **(E)** tumor xenografts. In each case, the left panel shows the T2-weighted MRI with the tumor region contoured in white while the right panel shows the corresponding lactate heatmap. **(F)** SNR of ^2^H-lactate in a 10.99 mm^3^ region of interest within the tumor or contralateral normal brain. Bars depict mean values and error bars represent standard deviation. * represents p<0.05, ** represents p<0.01, *** represents p<0.001 and **** represents p<0.0001.

### Lactate production from [U-^2^H]-pyruvate is an early biomarker of response to therapy in preclinical tumor models *in vivo*

We then examined the ability of [U-^2^H]-pyruvate to report on response to treatment with a pharmacological TERT inhibitor *in vivo*. 6-thio-dG is an analog of the TERT substrate 6-thio-dGTP that induces telomere uncapping in TERT+ cancer cells and is in clinical trials for solid tumors^9,10^. We treated mice bearing orthotopic glioblastoma (GBM6) or oligodendroglioma (BT88) tumor xenografts with 6-thio-dG and examined [U-^2^H]-pyruvate metabolism *in vivo* before (day 0) and after (day 6±1) treatment. Serial assessment of tumor volume by T2-weighted MRI indicated that 6-thio-dG induced tumor shrinkage, an effect that was significant starting day 14±1 for both models (see representative MRI and quantification in Fig. 7A-7B). Importantly, lactate production from [U-^2^H]-pyruvate was significantly reduced at an at an early time-point (day 7) at which no alteration in tumor volume could be detected by T2-weighted MRI (Fig. 7C-7D), suggesting that [U-^2^H]-pyruvate can report on response to pharmacological TERT inhibition *in vivo*.

**Figure 7.**
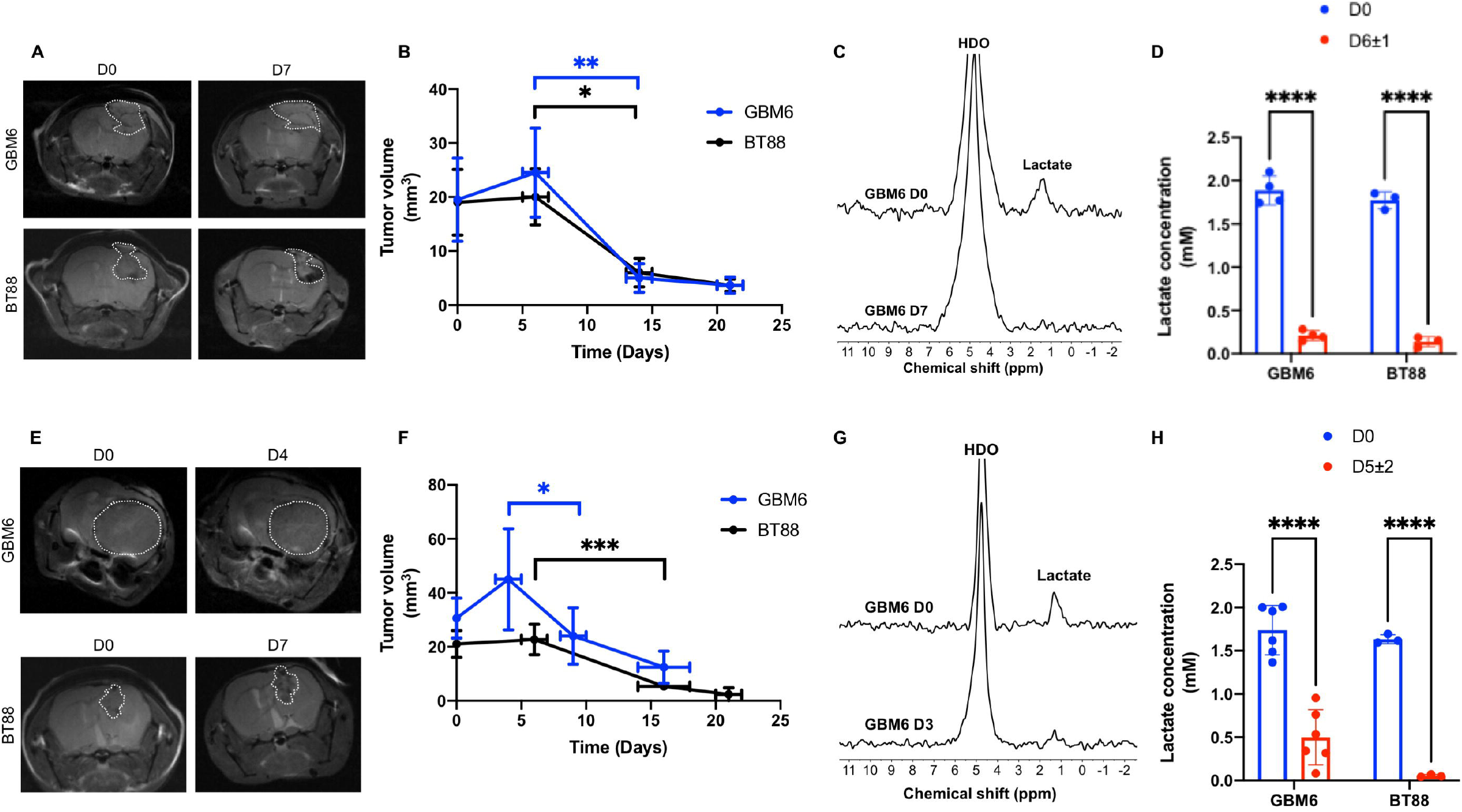
Lactate production from [U-^2^H]-pyruvate is an early biomarker of response to therapy *in vivo*. **(A)** Representative T2-weighted MRI of mice bearing orthotopic GBM6 (top row) or BT88 (bottom row) tumors before (day 0) and after (day 7) treatment with 6-thio-dG. **(B)** Effect of 6-thio-dG on tumor volume over time in mice bearing orthotopic GBM6 or BT88 tumor xenografts. **(C)** Representative spectra of [U-^2^H]-pyruvate metabolism from a mouse bearing an orthotopic GBM6 glioblastoma tumor before (day 0, top) and after (day 7, bottom) 6-thio-dG treatment. **(D)** Quantification of ^2^H-lactate production from [U-^2^H]-pyruvate before (day 0) and after (day 6±1) treatment with 6-thio-dG in mice bearing orthotopic GBM6 or BT88 tumors. **(E)** Representative T2-weighted MRI of mice bearing orthotopic GBM6 (top row) or BT88 (bottom row) tumors before (day 0) and after (day 4 or day 7 respectively) treatment with TMZ. **(F)** Effect of TMZ on tumor volume over time in mice bearing orthotopic GBM6 or BT88 tumor xenografts. **(G)** Representative spectra of [U-^2^H]-pyruvate metabolism from a mouse bearing an orthotopic GBM6 glioblastoma tumor before (day 0, top) and after (day 3, bottom) TMZ treatment. **(H)** Quantification of ^2^H-lactate production from [U-^2^H]-pyruvate before (day 0) and after (day 5±2) treatment with TMZ in mice bearing orthotopic GBM6 or BT88 tumors. Bars depict mean values and error bars represent standard deviation. * represents p<0.05, ** represents p<0.01, *** represents p<0.001 and **** represents p<0.0001.

Due to its essential role in maintaining replicative immortality, TERT is a biomarker of tumor proliferation and imaging biomarkers of TERT have the potential to report on response to cytocidal therapeutic agents. We, therefore, examined whether [U-^2^H]-pyruvate reports on response to temozolomide (TMZ), which is an alkylating chemotherapeutic drug that is standard of care for glioma patients^38,39^. We examined the effect of TMZ on tumor volume by T2-weighted MRI and on [U-^2^H]-pyruvate metabolism in mice bearing orthotopic glioblastoma (GBM6) or oligodendroglioma (BT88) tumor xenografts. As shown in Fig. 7E-7F, TMZ induced tumor shrinkage *in vivo*, an effect that was visible at day 9±1 in the GBM6 model and day 16±2 in the BT88 model. Importantly, [U-^2^H]-pyruvate metabolism to lactate was significantly reduced at day 5±2 in both GBM6 and BT88 models following TMZ treatment (Fig. 7G-7H) prior to tumor shrinkage. Collectively, these results suggest that lactate production from [U-^2^H]-pyruvate has the potential to serve as an early *in vivo* biomarker of response to therapy, including to targeted TERT inhibitors and standard chemotherapy.

## DISCUSSION

TERT expression is essential for tumor immortality in ∼85% of all human cancers^1,2^. In this study, we show that TERT acts via FOXO1 and NAMPT to upregulate NADH levels in multiple TERT-dependent cancers, including glioblastoma, oligodendroglioma, melanoma, neuroblastoma, and hepatocellular carcinoma. Importantly, we show that TERT-linked elevation in NADH is associated with elevated pyruvate flux to lactate, an effect which can be exploited for non-invasive ^2^H-MR metabolic imaging using [U-^2^H]-pyruvate (see schematic illustration in Fig. 2A). Our studies provide a way of integrating a mechanistic understanding of TERT-associated metabolic reprogramming with innovative metabolic imaging that has the potential to improve assessment of tumor burden and treatment response for cancer patients.

To the best of our knowledge for the first time, our study identifies a pan-cancer role for TERT in reprogramming tumor metabolism via the transcription factor FOXO1. Our results indicate that TERT expression is associated with higher inhibitory phosphorylation of FOXO1 and concomitantly reduced FOXO1 transcription factor activity in our tumor models. These results are in line with previous studies showing that FOXO1 phosphorylation leads to cytoplasmic sequestration that precludes activation of FOXO1 target genes in the nucleus^31^. Importantly, our study also identifies FOXO1 as a negative regulator of NAMPT, which is the rate-limiting enzyme in NADH biosynthesis^28^. Expression of a constitutively active form of FOXO1^32^ mimics the effects of TERT silencing and results in downregulation of NAMPT and concomitant loss of NADH. TERT expression is, therefore, associated with upregulation of NAMPT and elevated steady-state levels of NADH in a FOXO1-dependent manner in multiple patient-derived tumor models. Although the precise mechanism by which TERT mediates FOXO1 phosphorylation remains to be delineated, our studies highlight a previously unrecognized TERT-FOXO1 axis in the reprogramming of redox metabolism in cancer.

We have, for the first time, validated the utility of [U-^2^H]-pyruvate as an *in vivo* metabolic imaging agent and, importantly, demonstrated that lactate production from [U-^2^H]-pyruvate is linked to a critical oncogenic event i.e., TERT expression. Direct *in vivo* ^1^H-MRS-based detection of TERT-linked NAD(H) (representing the sum of NAD+ and NADH since these cannot be spectrally resolved) is complicated by issues of water suppression and the need for highly specialized acquisition schemes^40^. Lactate production from pyruvate is directly dependent on NADH and has long been used as a surrogate method for interrogating NADH levels in cell extracts^33,34^. Our studies indicate that [U-^2^H]-pyruvate flux to lactate is mechanistically linked to the TERT-mediated increase in NADH in cell and *in vivo* tumor models. ^2^H-MRS recently emerged as a novel method of imaging the metabolism of ^2^H-labeled substrates *in vivo* and studies have largely focused on assessing ^2^H-glucose metabolism in normal rat brain and in tumor-bearing animals^22,23,25-27^. Importantly, De Feyter et al. have established the clinical feasibility of using ^2^H-glucose to monitor tumor burden *in vivo* in glioblastoma patients^23^. Our studies extend the utility of ^2^H-MRS by leveraging a hitherto unexplored probe, i.e. [U-^2^H]-pyruvate for monitoring TERT expression and by demonstrating the ability of [U-^2^H]-pyruvate to assess tumor burden and response to therapy in a variety of preclinical tumor models *in vivo*.

Previous ^13^C-MRS studies have demonstrated that lactate production from hyperpolarized [1-^13^C]-pyruvate can be used for tumor imaging and treatment response assessment in preclinical cancer models^41,42^. The success of hyperpolarized [1-^13^C]-pyruvate underscores the utility of interrogating metabolic fluxes and has led to the initiation of clinical trials in several solid tumors. However, hyperpolarized ^13^C imaging is limited by the short lifetime of hyperpolarization and the requirement for access to expensive polarizers^41,42^. In this context, it should be noted that ^2^H-MRS following administration of [U-^2^H]-pyruvate provides a lower cost, technically less complicated and easy-to-implement method of interrogating pyruvate flux to lactate^22^. Importantly, [U-^2^H]-pyruvate is a safe, endogenous agent that has the properties needed for clinical application. First, while the concentration of [U-^2^H]-pyruvate used in our experiments (450 mg/kg) is 2-4 times higher than the concentration of hyperpolarized [1-^13^C]-pyruvate used in preclinical cancer models, it is similar to concentrations used in prior preclinical studies^43^ and did not lead to any adverse events in our studies. Studies with hyperpolarized [1-^13^C]-pyruvate have also established the safety of intravenous pyruvate administration in patients^41,44^. Second, although our studies were conducted at the higher field strength of 14.1T, the chemical shift difference between the HDO (4.75 ppm) and lactate (1.3 ppm) resonances allows complete spectral resolution at the clinically relevant field strength of 4T as demonstrated in previous studies with [6,6’-^2^H]-glucose in patients^23^. Third, the ability to cross the blood-brain barrier is an important consideration for imaging brain tumors, especially non-contrast enhancing low-grade gliomas^42^. Our studies indicate that [U-^2^H]-pyruvate is capable of imaging tumor burden and treatment response in patient-derived low-grade oligodendroglioma models. In essence, ^2^H-MRS in combination with [U-^2^H]-pyruvate has the potential to become a versatile, broadly applicable tool for clinical cancer metabolic imaging.

Our study highlights the ability of [U-^2^H]-pyruvate to provide an early readout of response to therapy *in vivo*. At present, response assessment in solid tumors is based on changes in tumor volume, which can take considerable time to manifest, thereby creating a need for early biomarkers of treatment response^12,13^. Using tumor models in which TERT expression is silenced in a doxycycline-inducible manner, we show that loss of TERT leads to reduced lactate production from [U-^2^H]-pyruvate. We also demonstrate that pharmacological inhibition of TERT using 6-thio-dG or chemotherapeutic inhibition of tumor proliferation using TMZ is accompanied by reduced [U-^2^H]-pyruvate flux to lactate. Strikingly, reduced lactate production is observed at early timepoints at which alterations in tumor volume cannot be observed by anatomical MR imaging. [U-^2^H]-pyruvate, therefore, has the potential to serve as a companion imaging agent for assessment of response to standard and emerging anti-cancer therapies, thereby aiding in their clinical translation and deployment.

In summary, our study has elucidated a novel mechanism of TERT-induced metabolic reprogramming in cancer. By inhibiting FOXO1, which functions as a negative regulator of NAMPT, TERT upregulates NADH biosynthesis. Importantly, ^2^H-MRS of [U-^2^H]-pyruvate metabolism can be used to monitor TERT expression *in vivo* and to provide a readout of response to therapy, prior to MRI-detectable changes in tumor volume. Clinical translation of our results has the potential to enable non-invasive annotation of TERT expression *in vivo* and to aid in the assessment of response to anti-cancer therapies.

## METHODS

### Cell culture

GBM1 and GBM6 were isolated from isocitrate dehydrogenase wild-type glioblastoma patients as previously described^3,8,45^. SF10417 and BT88 cells were isolated from patients harboring isocitrate dehydrogenase mutant oligodendrogliomas^46,47^. A375, HepG2 and SK-N-SH cells were a kind gift from Dr. Joseph Costello and have been previously described^3^. GBM1 cells were maintained as monolayers in DMEM/Ham’s F-12 1:1 media supplemented with 2 mM glutamine, 10% fetal calf serum and 1% penicillin/streptomycin^3,8^. SF10417 cells were maintained as monolayers in laminin-coated flasks and were cultured in serum-free Neurocult NS-A media containing 2 mM glutamine, 1% penicillin/streptomycin, B-27 and N2 supplements, 20 ng/mL EGF, 20 ng/mL FGF and 20 ng/mL PDGF-AA^46^. GBM6, BT88, A375, HepG2 and SK-N-SH cells were maintained as monolayers in DMEM supplemented with 2 mM glutamine, 10% fetal calf serum and 1% penicillin/streptomycin^3,45,47^. All cell lines were routinely tested for mycoplasma contamination, authenticated by short tandem repeat fingerprinting (Cell Line Genetics) and assayed within 6 months of authentication.

For exogenous expression of CA-FOXO1, TERT+ cells were transiently transfected with human CA-FOXO1 (pcDNA3 Flag FKHR AAA mutant; gift from Kunliang Guan; Addgene plasmid # 13508) using polyethyleneimine. This plasmid encodes a FLAG-tagged FOXO1 protein in which all 3 phosphorylation sites have been mutated to alanine (T24A, S256A, and S319A), thereby leading to expression of constitutively active and nuclear FOXO1. CA-FOXO1 expression was verified at 72 h post-transfection by western blotting for the FLAG tag.

For doxycycline-inducible TERT silencing, microRNA-embedded miR-E shRNA against TERT (TGCTGTTGACAGTGAGCGATCAGGTCTTTCTTTTATGTCATAGTGAAGCCACAGATGTATGACATAAAAGAAAGACCTGAGTGCCTACTGCCTCGGA) was designed using Splash RNA and cloned into a custom-designed lentiviral vector (N174-CMV-TurboGFP-miR-E-IRES-puro). This vector was generated by modifying the constitutive lentiviral vector, N174-MCS-puro (Addgene #81068) through a sequential cloning process to contain a single CMV promoter driving expression of the TurboGFP-miR-E-IRES-puro transcript. Lentiviral particles were produced by transient transfection of psPAX2 (Addgene #12260), pMD2.G (Addgene #12259), and transfer plasmid (N174-CMV-GFP-miR-E-IRES-puro-shRNA) into HEK293T cells with Lipofectamine 2000 reagent (Life Technologies) according to the manufacturer’s instructions. HepG2 cells were transfected with lentivirus using polybrene followed by puromycin selection.

### Quantitative RT-PCR

Gene expression was measured by quantitative RT-PCR and normalized to β-actin^8,48^ The SYBR Green quantitative RT-PCR kit (Sigma) was used with the following primers: TERT (forward primer: TCACGGAGACCACGTTTCAAA; reverse primer: TTCAAGTGCTGTCTGATTCCAAT), NAMPT (forward primer: GTTCCAGCAGCAGAACACAG; reverse primer: GCTGACCACAGATACAGGCA), FOXO1 (forward primer: GCAGCCAGGCATCTCATAA; reverse primer: CCTACCATAGCCATTGCAGC) and β-actin (forward primer: AGAGCTACGAGCTGCCTGAC; reverse primer: AGCACTGTGTTGGCGTACAG).

### Metabolite and Activity assays

Telomerase activity was confirmed using the TRAPeze® RT kit (Sigma) according to manufacturer’s instructions. NAMPT activity was measured using a kit (Abcam, catalog # ab221819). FOXO1 transcription factor activity assay was measured using a commercially available kit (Abcam, catalog # ab207204), which provides an ELISA-based method of measuring FOXO1 activation in extracts. Briefly, the assay measures binding of active FOXO1 to a DNA sequence containing the FOXO1 consensus binding site that is immobilized on a 96-well plate. FOXO1 detection is achieved using a primary antibody that recognizes an epitope of FOXO1 accessible only when the protein is activated and bound to its target DNA. Levels of NAD+ and NADH were measured in cell extracts using a kit (Biovision, catalog # K337). For supplementation studies, NA (1 mM) or tryptophan (0.25 mM) was added to the medium prior to extraction for NAD+ and NADH measurement. Levels of GSH and GSSG were measured using the GSH/GSSG Ratio Detection Assay Kit II (Abcam catalog # ab205811). NADPH and NADP+ were measured using the NADP/NADPH assay kit (Abcam catalog # 176724).

### RNA interference

SMARTpool siRNAs (Dharmacon) against human TERT (M-003547-02) and non-targeting siRNA pool #2 (D-001206-14-05) were used to transiently silence gene expression for 72 h as described previously^49^.

### Western blotting

Cells (∼5 × 10^6^) were lysed by sonication in RIPA buffer (25 mM Tris-HCl pH 7.6, 150 mM NaCl, 1% NP-40, 1% sodium deoxycholate, 0.1% SDS) containing 150 nM aprotinin, 1 µM each of leupeptin and E64 protease inhibitor and phosphatase inhibitor cocktail (Sigma, catalog# P5726). For FOXO-1 analysis, nuclear extracts were prepared using the NE-PER fractionation kit (Thermo-Fisher Scientific) according to manufacturer’s instructions. Lysates were cleared by centrifugation at 14,000 rpm for 15 minutes at 4 °C and boiled in SDS-PAGE sample buffer (95°C for 10 minutes). Protein (∼ 20 µg) was separated on a 10% polyacrylamide gel (Bio-Rad) by sodium dodecyl sulphate polyacrylamide gel electrophoresis, transferred onto Immobilon-FL PVDF membrane (Millipore) and probed for FOXO1 (Cell Signaling, #2880), phospho-FOXO1 S256 (Cell Signaling, #84192), FLAG (Cell Signaling, anti-FLAG M2, # 14793). β-Actin (Cell Signaling, 4970) was used as loading control.

### ^2^H-MRS of live cells

Cells were incubated in media containing 10 mM [U-^2^H]-pyruvate for 72 h. Live cells were harvested, suspended in saline in 12 mm glass vials and ^2^H-MR spectra acquired using a 16 mm ^2^H single loop surface coil (DOTY Scientific) at a Varian 14.1T vertical MR scanner (Agilent Technologies). A pulse-acquire sequence (TR=260ms, NA=2500, complex points=512, flip angle=64°, spectral width=2kHz) was used. Corrected amplitudes (for saturation) of fitted water and lactate peaks were converted to concentration in millimolar using the natural abundance HDO signal (12.8 mM) collected from a similar vial containing only saline. The latter was determined assuming a 55.5 M water concentration and a deuterium natural abundance of 0.0115%. Data analysis was performed using MestReNova (v12.0.4, Mestrelab, Spain).

### Orthotopic tumor xenograft generation and MRI

Animal studies were conducted in accordance with UCSF Institutional Animal Care and Use Committee (IACUC) guidelines. GBM1, GBM6, BT88 or SF10417 cells (2×10^5^ cells for SF10417 and 3×10^5^ for all other cell lines in 3µl saline solution) were intracranially injected into SCID mice (female, FOX Chase C.B.17 SCID, 5-6-weeks old) by the free-hand technique^49-51^. HepG2_dox-TERT_ cells (1×10^6^ in 100µl saline) were injected subcutaneously into SCID mice^52^. Once tumors reached a volume of 27.0 ± 7.8 mm^3^ (orthotopic brain) or 180.3 ± 24.7 mm^3^ (subcutaneous), this timepoint was considered day zero (D0). Mice were randomized and treated with vehicle-control (saline), TMZ (50 mg/kg), 6-thio-dG (50 mg/kg) or doxycycline (50 mg/kg) daily via intraperitoneal injection. Animals under doxycycline treatment also received *ad libitum* access to doxycycline enriched food. In total, 44 tumor bearing and 3 healthy (tumor-free controls) mice were investigated. All studies were performed on a Varian 14.1T vertical MR scanner (Agilent Technologies) equipped with a single-channel ^1^H volume coil. Animals were anesthetized and maintained using isoflurane (1-2% in O_2_) and placed headfirst in the prone position with a respiratory sensor. Axial anatomical T2-weighted or T1-weighted images were acquired using either a spin echo multi-slice sequence (TE/TR=20/1200 ms, FOV=30×30 mm^2^, matrix=256×256, slice thickness=1 mm, NA=4)^49-51^ or a gradient echo multi-slice sequence (TE/TR=2.09/120 ms, FOV=30×30 mm^2^, matrix=256×256, slice thickness=1 mm, NA=10) respectively. Tumor contours in each axial slice were drawn manually and tumor volume was evaluated as the sum of the areas multiplied by slice thickness using in-house software^49-51^.

### ^2^H-MRS *in vivo*

Studies were performed on a Varian 14.1T vertical MR scanner using a 16 mm ^2^H single loop surface coil (DOTY Scientific). The coil loop was placed on top of the animal head or the subcutaneous tumor. Following intravenous injection of a bolus of [U-^2^H]-pyruvate in isotonic buffer (Sigma-Aldrich, 450 mg/kg dissolved in 40 mM Tris-HCl, 0.3 mM EDTA, pH 7) over 1 minute, non-localized ^2^H-MR spectra were acquired over 16 minutes with a pulse-acquire sequence (TR=500 ms, NA=500, complex points=512, flip angle=64, spectral width=2 kHz, temporal resolution = 4 min 10 s). Non-localized ^2^H-MR spectra were analyzed using MestReNova. Metabolites were referenced to semi-heavy water (HDO; 4.75 ppm). Absolute lactate concentrations were determined by correcting peak integrals for saturation effects and normalizing to pre-injection HDO (estimated to be 10.12 mM as described in previous studies^23,25,26^).

For spatial localization, a 2D chemical shift imaging (CSI) sequence with a temporal resolution of 4 min 10 sec (TE/TR=1.35/250 ms, FOV=30×30×8 mm^3^, 128 points, 2.5 kHz spectral width, NA=20) resulting in a nominal voxel size of 112.5 µl was used. The data were analyzed using adapted in-house Matlab codes (R2021a, Mathworks)^18,19^. For each voxel at every time point, spectra were analyzed after a 10 Hz line broadening by determining the area under each peak by integration. For generation of [3,3’-^2^H]-lactate SNR heatmaps, raw data were interpolated from 8 × 8 matrix to a 256 × 256 matrix using the *imresize* function of Matlab and the Lanczos-2 interpolation algorithm and normalizing to noise, which was evaluated as the standard deviation of the real part of the signal in a voxel outside of the brain. The SNR of [3,3’-^2^H]-lactate was assessed in a 10.99 mm^3^ volume from tumor and contralateral normal brain.

### Statistical analysis

All experiments were performed on a minimum of 3 samples (n≥3) and results presented as mean ± standard deviation. Statistical significance was assessed in GraphPad Prism 9 using a two-way ANOVA, or two-tailed Student’s T-test assuming unequal variance with p<0.05 considered significant. * represents p<0.05, ** represents p<0.01, *** represents p<0.001 and **** represents p<0.0001.

### Data availability

Data included in this manuscript that support the findings and conclusions of this study are available from the corresponding author upon reasonable request.

## ACKNOWLEDGEMENTS

This work was supported by the National Institutes of Health (R01CA239288, R01CA172845, R01CA197254, R01NS105087, P01CA118816,) Department of Defense (W81XWH201055315), UCSF Brain Tumor Center Loglio Collective and NICO.

## AUTHOR CONTRIBUTIONS

P.V. conceptualized the research and directed the studies; P.V., G.B., C.T., N.S., C.B., and M.T. performed and analyzed the experiments; A.M.G. assisted with cell and *in vivo* studies; J.F.C provided cell lines and reagents; P.V. and G.B. wrote the manuscript; P.V. and S.M.R. secured funding.

## COMPETING FINANCIAL INTERESTS

The authors declare no competing financial interests.

## SUPPLEMENTARY FIGURE LEGENDS

**Supplementary Figure 1.**
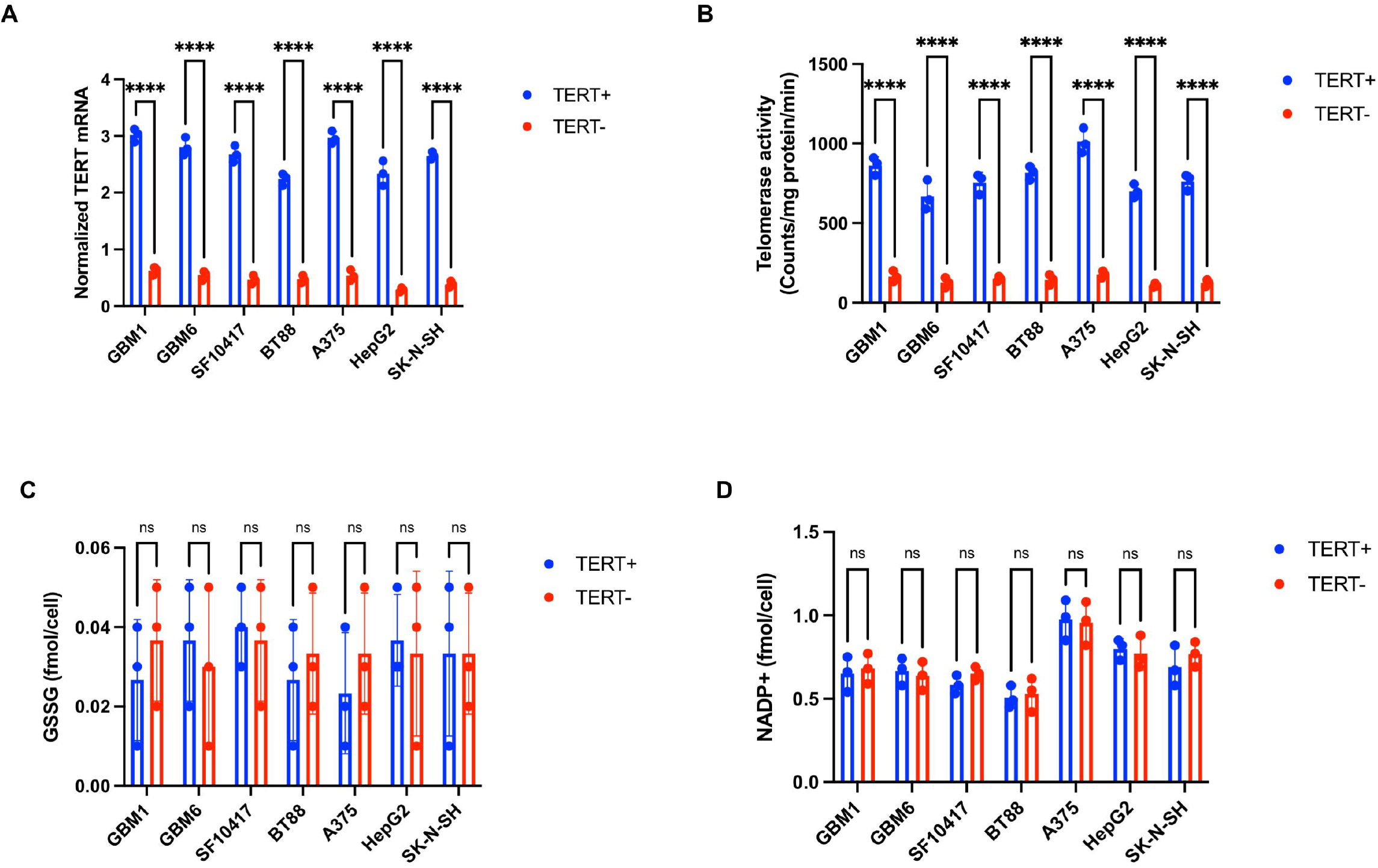
Effect of TERT silencing on tumor metabolism. Normalized TERT mRNA **(A)**, telomerase activity **(B)**, GSSG **(C)** and NADP^+^ **(D)** in TERT+ and TERT-glioblastoma (GBM1, GBM6), oligodendroglioma (SF10417, BT88), melanoma (A375), hepatocellular carcinoma (HepG2) and neuroblastoma (SK-N-SH) cells. Bars depict mean values and error bars represent standard deviation. * represents p<0.05, ** represents p<0.01, *** represents p<0.001 and **** represents p<0.0001.

**Supplementary Figure 2.**
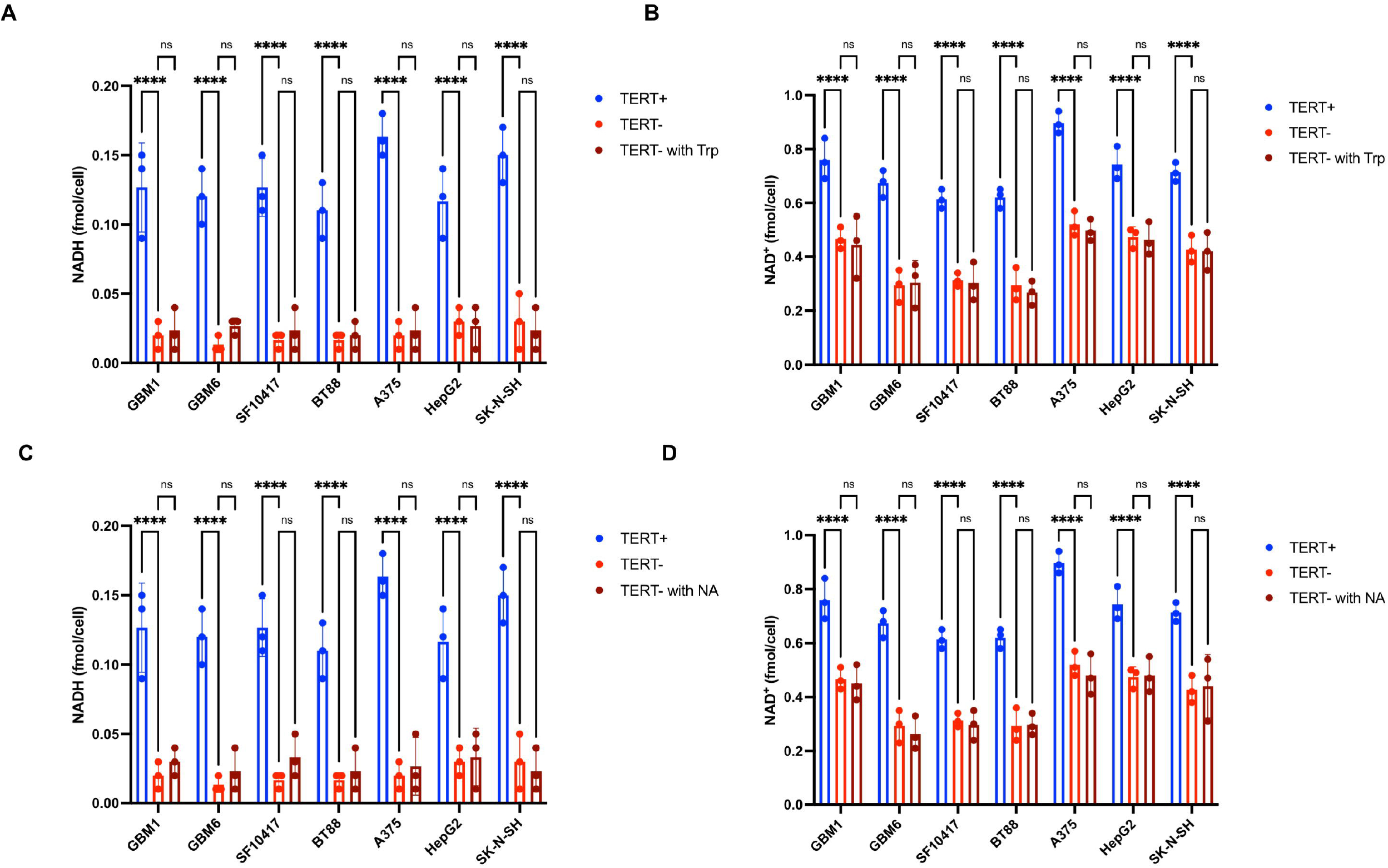
Supplementation with tryptophan or nicotinic acid does not rescue NAD^+^ and NADH levels in TERT-cells. Levels of NADH **(A)** and NAD+ **(B)** in TERT+ cells, TERT-cells and TERT-cells supplemented with tryptophan in glioblastoma (GBM1, GBM6), oligodendroglioma (SF10417, BT88), melanoma (A375), hepatocellular carcinoma (HepG2) and neuroblastoma (SK-N-SH) models. Levels of NADH **(C)** and NAD+ **(D)** in TERT+ cells, TERT-cells and TERT-cells supplemented with nicotinic acid in glioblastoma (GBM1, GBM6), oligodendroglioma (SF10417, BT88), melanoma (A375), hepatocellular carcinoma (HepG2) and neuroblastoma (SK-N-SH) models. Bars depict mean values and error bars represent standard deviation. * represents p<0.05, ** represents p<0.01, *** represents p<0.001 and **** represents p<0.0001.

**Supplementary Figure 3.**
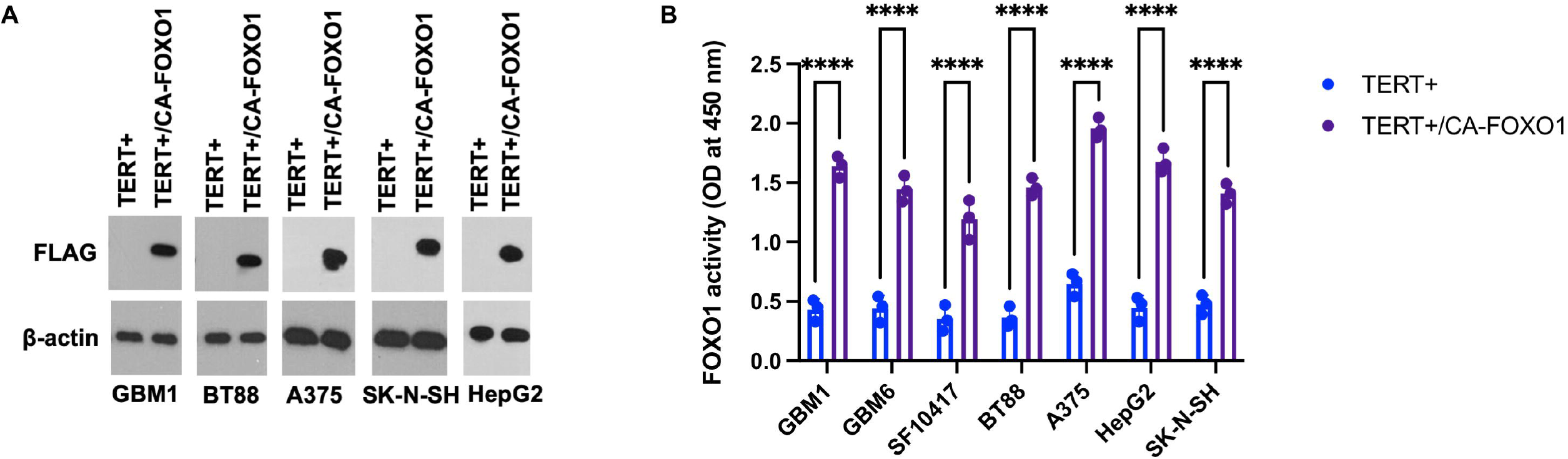
Validation of expression of a constitutively active form of FOXO1 (CA-FOXO1) in TERT+ cells. **(A)** Western blotting for the FLAG tag in TERT+ cells with and without exogenous expression of CA-FOXO1 in glioblastoma (GBM1), oligodendroglioma (BT88), melanoma (A375), neuroblastoma (SK-N-SH) and hepatocellular carcinoma (HepG2) models. β-actin was used as loading control. **(B)** FOXO1 transcription factor activity in TERT+ cells with and without exogenous expression of CA-FOXO1 in glioblastoma (GBM1, GBM6), oligodendroglioma (SF10417, BT88), melanoma (A375), neuroblastoma (SK-N-SH) and hepatocellular carcinoma (HepG2) models. Bars depict mean values and error bars represent standard deviation. * represents p<0.05, ** represents p<0.01, *** represents p<0.001 and **** represents p<0.0001.

**Supplementary Figure 4.**
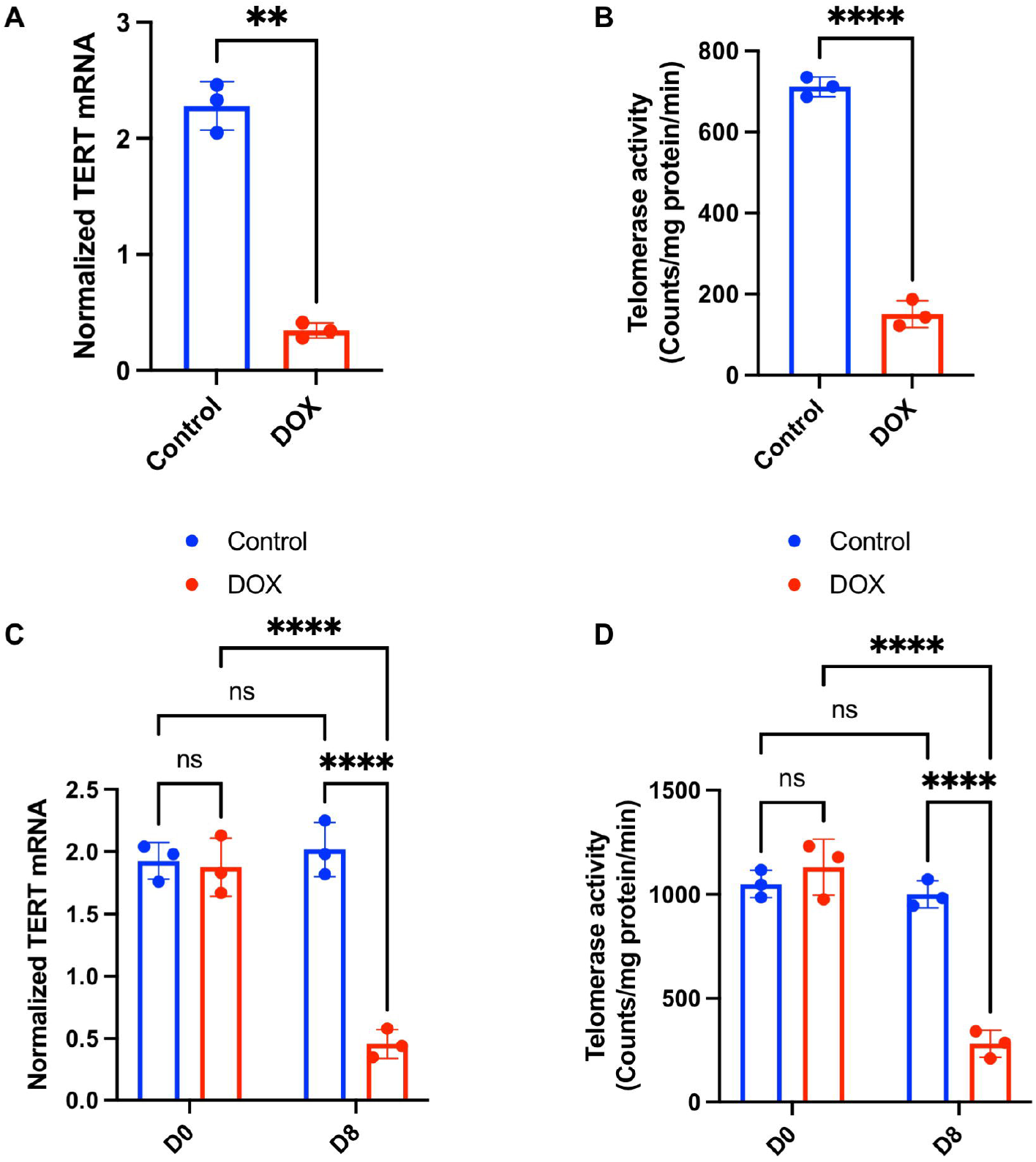
Validation of doxycycline-inducible TERT silencing in HepG2_dox-TERT_ cells and tumors. Normalized TERT mRNA **(A)** and telomerase activity **(B)** in HepG2_dox-TERT_ cells treated with vehicle-control (saline) or doxycycline for 72 h. Quantification of TERT mRNA **(C)** and telomerase activity **(D)** before (day 0) and after (day 8) treatment with vehicle-control (saline) or doxycycline in mice bearing subcutaneous HepG2_dox-TERT_ tumors. Bars depict mean values and error bars represent standard deviation. * represents p<0.05, ** represents p<0.01, *** represents p<0.001 and **** represents p<0.0001.

## Notes

### Competing Interest Statement

The authors have declared no competing interest.

## REFERENCES

1 Shay, J. W. & Wright, W. E. Telomeres and telomerase: three decades of progress. Nature Reviews Genetics 20, 299–309, doi:10.1038/s41576-019-0099-1 (2019).

2 Bell, R. J. et al. Understanding TERT Promoter Mutations: A Common Path to Immortality. Molecular cancer research: MCR 14, 315–323, doi:10.1158/1541-7786.mcr-16-0003 (2016).

3 Bell, R. J. et al. Cancer. The transcription factor GABP selectively binds and activates the mutant TERT promoter in cancer. Science 348, 1036–1039, doi:10.1126/science.aab0015 (2015).

4 Akincilar, S. C. et al. Long-Range Chromatin Interactions Drive Mutant TERT Promoter Activation. Cancer discovery 6, 1276–1291, doi:10.1158/2159-8290.Cd-16-0177 (2016).

5 Korber, V. et al. Evolutionary Trajectories of IDH(WT) Glioblastomas Reveal a Common Path of Early Tumorigenesis Instigated Years ahead of Initial Diagnosis. Cancer cell 35, 692–704 e612, doi:10.1016/j.ccell.2019.02.007 (2019).

6 Li, Y. et al. Non-canonical NF-kappaB signalling and ETS1/2 cooperatively drive C250T mutant TERT promoter activation. Nat Cell Biol 17, 1327–1338 (2015).

7 Chiba, K. et al. Cancer-associated TERT promoter mutations abrogate telomerase silencing. eLife 4, doi:10.7554/eLife.07918 (2015).

8 Mancini, A. et al. Disruption of the beta1L Isoform of GABP Reverses Glioblastoma Replicative Immortality in a TERT Promoter Mutation-Dependent Manner. Cancer cell 34, 513–528 e518 (2018).

9 Mender, I., Gryaznov, S., Dikmen, Z. G., Wright, W. E. & Shay, J. W. Induction of telomere dysfunction mediated by the telomerase substrate precursor 6-thio-2’-deoxyguanosine. Cancer discovery 5, 82–95, doi:10.1158/2159-8290.Cd-14-0609 (2015).

10 Sengupta, S. et al. Induced Telomere Damage to Treat Telomerase Expressing Therapy-Resistant Pediatric Brain Tumors. Molecular cancer therapeutics 17, 1504–1514, doi:10.1158/1535-7163.Mct-17-0792 (2018).

11 Kauppinen, R. A. & Peet, A. C. Using magnetic resonance imaging and spectroscopy in cancer diagnostics and monitoring: preclinical and clinical approaches. Cancer Biol Ther 12, 665–679, doi:10.4161/cbt.12.8.18137 (2011).

12 García-Figueiras, R. et al. How clinical imaging can assess cancer biology. Insights Imaging 10, 28, doi:10.1186/s13244-019-0703-0 (2019).

13 Brindle, K. New approaches for imaging tumour responses to treatment. Nature reviews. Cancer 8, 94–107, doi:10.1038/nrc2289 (2008).

14 Hygino da Cruz, L. C., Rodriguez, I., Domingues, R. C., Gasparetto, E. L. & Sorensen, A. G. Pseudoprogression and Pseudoresponse: Imaging Challenges in the Assessment of Posttreatment Glioma. American Journal of Neuroradiology 32, 1978, doi:10.3174/ajnr.A2397 (2011).

15 Zikou, A. et al. Radiation Necrosis, Pseudoprogression, Pseudoresponse, and Tumor Recurrence: Imaging Challenges for the Evaluation of Treated Gliomas. Contrast media & molecular imaging 2018, 6828396–6828396, doi:10.1155/2018/6828396 (2018).

16 Indran, I. R., Hande, M. P. & Pervaiz, S. hTERT overexpression alleviates intracellular ROS production, improves mitochondrial function, and inhibits ROS-mediated apoptosis in cancer cells. Cancer research 71, 266–276, doi:10.1158/0008-5472.can-10-1588 (2011).

17 Ahmad, F. et al. Nrf2-driven TERT regulates pentose phosphate pathway in glioblastoma. Cell death & disease 7, e2213, doi:10.1038/cddis.2016.117 (2016).

18 Viswanath, P. et al. Metabolic imaging detects elevated glucose flux through the pentose phosphate pathway associated with TERT expression in low-grade gliomas. Neuro-oncology, doi:10.1093/neuonc/noab093 (2021).

19 Viswanath, P. et al. Non-invasive assessment of telomere maintenance mechanisms in brain tumors. Nature communications 12, 92, doi:10.1038/s41467-020-20312-y (2021).

20 Glunde, K. & Bhujwalla, Z. M. Metabolic Tumor Imaging Using Magnetic Resonance Spectroscopy. Seminars in Oncology 38, 26–41, doi:10.1053/j.seminoncol.2010.11.001 (2011).

21 Ishii, N. et al. Multiple high-throughput analyses monitor the response of E. coli to perturbations. Science 316, 593–597, doi:10.1126/science.1132067 (2007).

22 De Feyter, H. M. & de Graaf, R. A. Deuterium metabolic imaging - Back to the future. Journal of magnetic resonance (San Diego, Calif.: 1997) 326, 106932, doi:10.1016/j.jmr.2021.106932 (2021).

23 De Feyter, H. M. et al. Deuterium metabolic imaging (DMI) for MRI-based 3D mapping of metabolism in vivo. Sci Adv 4, eaat7314 (2018).

24 Hesse, F. et al. Monitoring tumor cell death in murine tumor models using deuterium magnetic resonance spectroscopy and spectroscopic imaging. Proceedings of the National Academy of Sciences of the United States of America 118, doi:10.1073/pnas.2014631118 (2021).

25 Kreis, F. et al. Measuring Tumor Glycolytic Flux in Vivo by Using Fast Deuterium MRI. Radiology 294, 289–296, doi:10.1148/radiol.2019191242 (2020).

26 Lu, M., Zhu, X. H., Zhang, Y., Mateescu, G. & Chen, W. Quantitative assessment of brain glucose metabolic rates using in vivo deuterium magnetic resonance spectroscopy. Journal of cerebral blood flow and metabolism: official journal of the International Society of Cerebral Blood Flow and Metabolism 37, 3518–3530 (2017).

27 Markovic, S. et al. Deuterium MRSI characterizations of glucose metabolism in orthotopic pancreatic cancer mouse models. NMR in biomedicine 34, e4569, doi:10.1002/nbm.4569 (2021).

28 Garten, A., Petzold, S., Körner, A., Imai, S.-i. & Kiess, W. Nampt: linking NAD biology, metabolism and cancer. Trends in Endocrinology & Metabolism 20, 130–138, doi:10.1016/j.tem.2008.10.004 (2009).

29 Gorrini, C., Harris, I. S. & Mak, T. W. Modulation of oxidative stress as an anticancer strategy. Nat Rev Drug Discov 12, 931–947, doi:10.1038/nrd4002 (2013).

30 Xie, N. et al. NAD(+) metabolism: pathophysiologic mechanisms and therapeutic potential. Signal Transduct Target Ther 5, 227, doi:10.1038/s41392-020-00311-7 (2020).

31 Gross, D. N., van den Heuvel, A. P. J. & Birnbaum, M. J. The role of FoxO in the regulation of metabolism. Oncogene 27, 2320–2336, doi:10.1038/onc.2008.25 (2008).

32 Tang, E. D., Nuñez, G., Barr, F. G. & Guan, K. L. Negative regulation of the forkhead transcription factor FKHR by Akt. The Journal of biological chemistry 274, 16741–16746, doi:10.1074/jbc.274.24.16741 (1999).

33 Day, S. E. et al. Detecting tumor response to treatment using hyperpolarized 13C magnetic resonance imaging and spectroscopy. Nat Med 13, 1382–1387, doi:10.1038/nm1650 (2007).

34 Williamson, D. H., Lund, P. & Krebs, H. A. The redox state of free nicotinamide-adenine dinucleotide in the cytoplasm and mitochondria of rat liver. The Biochemical journal 103, 514–527, doi:10.1042/bj1030514 (1967).

35 Guarino, V. A., Oldham, W. M., Loscalzo, J. & Zhang, Y. Y. Reaction rate of pyruvate and hydrogen peroxide: assessing antioxidant capacity of pyruvate under biological conditions. Sci Rep 9, 19568, doi:10.1038/s41598-019-55951-9 (2019).

36 Arce-Molina, R. et al. A highly responsive pyruvate sensor reveals pathway-regulatory role of the mitochondrial pyruvate carrier MPC. eLife 9, doi:10.7554/eLife.53917 (2020).

37 Patel, P. L., Suram, A., Mirani, N., Bischof, O. & Herbig, U. Derepression of hTERT gene expression promotes escape from oncogene-induced cellular senescence. Proceedings of the National Academy of Sciences 113, E5024 (2016).

38 Weller, M. et al. EANO guidelines on the diagnosis and treatment of diffuse gliomas of adulthood. Nat Rev Clin Oncol, doi:10.1038/s41571-020-00447-z (2020).

39 Stupp, R. et al. Radiotherapy plus concomitant and adjuvant temozolomide for glioblastoma. The New England journal of medicine 352, 987–996, doi:10.1056/NEJMoa043330 (2005).

40 de Graaf, R. A. & Behar, K. L. Detection of cerebral NAD(+) by in vivo (1)H NMR spectroscopy. NMR in biomedicine 27, 802–809, doi:10.1002/nbm.3121 (2014).

41 Kurhanewicz, J. et al. Hyperpolarized (13)C MRI: Path to Clinical Translation in Oncology. Neoplasia (New York, N.Y.) 21, 1–16 (2019).

42 Viswanath, P., Li, Y. & Ronen, S. M. in Glioma Imaging: Physiologic, Metabolic, and Molecular Approaches (ed Whitney B. Pope) 191–209 (Springer International Publishing, 2020).

43 Suh, S. W., Aoyama, K., Matsumori, Y., Liu, J. & Swanson, R. A. Pyruvate administered after severe hypoglycemia reduces neuronal death and cognitive impairment. Diabetes 54, 1452–1458, doi:10.2337/diabetes.54.5.1452 (2005).

44 Nelson, S. J. et al. Metabolic imaging of patients with prostate cancer using hyperpolarized [1-(1)(3)C]pyruvate. Science translational medicine 5, 198ra108, doi:10.1126/scitranslmed.3006070 (2013).

45 Sarkaria, J. N. et al. Use of an orthotopic xenograft model for assessing the effect of epidermal growth factor receptor amplification on glioblastoma radiation response. Clinical cancer research: an official journal of the American Association for Cancer Research 12, 2264–2271, doi:10.1158/1078-0432.Ccr-05-2510 (2006).

46 Jones, L. E. et al. Patient-derived cells from recurrent tumors that model the evolution of IDH-mutant glioma. Neurooncol Adv 2, vdaa088 (2020).

47 Kelly, J. J. et al. Oligodendroglioma cell lines containing t(1;19)(q10;p10). Neurooncology 12, 745–755, doi:10.1093/neuonc/noq031 (2010).

48 Ohba, S. et al. Mutant IDH1 Expression Drives TERT Promoter Reactivation as Part of the Cellular Transformation Process. Cancer research 76, 6680–6689 (2016).

49 Viswanath, P. et al. 2-hydroxyglutarate-mediated autophagy of the endoplasmic reticulum leads to an unusual downregulation of phospholipid biosynthesis in mutant IDH1 gliomas. Cancer research, doi:10.1158/0008-5472.can-17-2926 (2018).

50 Batsios, G. et al. PI3K/mTOR inhibition of IDH1 mutant glioma leads to reduced 2HG production that is associated with increased survival. Sci Rep 9, 10521, doi:10.1038/s41598-019-47021-x (2019).

51 Viswanath, P. et al. Mutant IDH1 gliomas downregulate phosphocholine and phosphoethanolamine synthesis in a 2-hydroxyglutarate-dependent manner. Cancer & metabolism 6, 3, doi:10.1186/s40170-018-0178-3 (2018).

52 Schnater, J. M. et al. Subcutaneous and intrahepatic growth of human hepatoblastoma in immunodeficient mice. Journal of hepatology 45, 377–386, doi:10.1016/j.jhep.2006.03.018 (2006).

